# Integrated analysis and annotation for T-cell receptor sequences using TCRosetta

**DOI:** 10.1101/2023.02.20.529199

**Authors:** Tao Yue, Si-Yi Chen, Wen-Kang Shen, Liming Cheng, An-Yuan Guo

## Abstract

**Background:** T cells and T cell receptors (TCRs) are essential components of the adaptive immune system. TCRs, on the surface of T cells, are responsible for recognizing and binding antigen peptide-MHC complex and play vital roles in T-cell immunology. Characterization of TCR repertoire offers a promising and high informative source for understanding the functions of T cells in immune responses and immunotherapies. Many researchers are now interested in TCR repertoire study; however, there are rare online servers for TCR analysis, especially no servers for TCR annotation and advanced analyses.

**Results:** We developed TCRosetta, a comprehensive online server integrating the analytical methods for TCR repertoire/sequences analysis and visualization. TCRosetta combines general features analysis, large-scale sequence clustering, network construction, TCR-peptide binding prediction, generation probability calculation, and k-mer motif analysis for robust TCR sequence analysis, making TCR data analysis as simple as possible and allowing users to concentrate on research rather than coding. In addition, TCRosetta could annotate disease information for TRB CDR3 sequences by batch searching similar sequences in manually curated disease-related TCR database. The TCRosetta server accepts multiple input data formats and can analyze ∼20000 TCR sequences in less than three minutes.

**Conclusions:** TCRosetta is the most comprehensive web server to date for TCR sequences/repertoires analysis and it is freely available at http://bioinfo.life.hust.edu.cn/TCRosetta/. It can be applied to help discover novel biomarkers for disease diagnosis and identify cancer-associated TCR sequences.

**WHAT IS ALERADY KNOWN ON THIS TOPIC:** T cell receptor repertoires are largely untapped resource than can be used for predicting immune responses to different exposures including viral infections and tumor neoantigens. The downstream analysis of TCR repertoire is often performed by different tools requiring diverse operating environments and expertise. There is no webserver for comprehensive TCR repertoire analysis including general and advanced analysis.

**WHAT THIS STUDY ADDS:** We developed TCRosetta, a comprehensive platform for analyzing T-cell repertoire which combines nearly all TCR analysis methods. It supports different kinds of the format of input including most mainstream TCR extraction tools or amino acid sequences. It could analyze the features of TCR repertoire and display them in interactive graphs and is the first platform with a batch search and TCR annotation function.

**HOW THIS STUDY MIGHT AFFECT RESEARCH, PACTICE OR POLICY:** TCRosetta can be applied to discover novel biomarkers to predict response in immunotherapy such TCR repertoire diversity and clonality. It also can identify cancer-associated TCR sequences by clustering biochemically similar CDR3 sequences. It can make TCR repertoire analysis as effortless as possible and help users focus on research instead of coding.

## Background

T-cell receptor (TCR) is the T-cell surface protein complex that identifies antigenic peptides bound to major histocompatibility complex molecules (MHC). A TCR is a heterodimer composed of two chains (αβ or γδ), and the number of different TCRs in the human body is estimated to vary from 10^12^ to 10^18^ [1]. The high diversity of TCR is the result of genetic rearrangement of the variable (V), diversity (D), and joining (J) genes, commonly known as V(D)J recombination, a mechanism of somatic recombination that occurs during the maturation of T-cell [2]. Complementary-determining region 3 (CDR3) is the most variable component of the TCR, which directly interacts with epitope and is regarded as the main driver of T-cell specificity [3].

In recent years, increased emphasis has been placed on the downstream analysis of TCR data, which can be roughly categorized into two major sections: general and advanced analyses [4]. The general analyses include diversity, clonality, CDR3 length distribution, clonal composition, V/J gene usage, V-J gene utilization. Analyzing general features of the TCR repertoire can contribute to an understanding of immune response [5]. For example, the diversity of the TCR repertoire is considered a potential biomarker for tracking the response to immunotherapy [6]. Features of the TCRβ chain (TRB) can also assist cancer early-stage diagnosis [7] and treatment selection [8]. Besides, the advanced analyses include network analysis, clustering, enrichment analysis and the prediction of TCR-peptide binding, which support further clinical development of immunotherapy. For example, clustering similar TCR sequence to disease-specific TCR sequences could help to make TCR-engineered T-cells specifically recognize tumor antigens [9].

Until now, some tools for analyzing and visualizing general features of the TCR repertoire have been developed, such as tcR [10], VDJtools [11], and VisTCR [12]. There are also various methods for advanced analyses of TCR repertoire. TCRdist [13], GLIPH [14], and GIANA [15] all focus on finding antigen-specific TCR based on sequence similarity. IGoR [16] and OLGA [17] could compute TCR amino acid sequence generation probabilities. However, these methods or tools can only be run locally in different environments and are challenging for users without programming knowledge. Although some online platforms for analyzing TCR repertoire have been developed for, they all need registration and can only analyze the repertoire’s general features. Therefore, it is essential to provide a user-friendly online server for comprehensive TCR sequence analysis and provide more valuable information. In this study, we introduce TCRosetta (http://bioinfo.life.hust.edu.cn/TCRosetta/), a powerful and comprehensive server for analyzing and visualizing the TCR sequence.

## Methods

### Data pre-processing

All input TCR sequences will be subjected to quality control to assure their dependability and quality, based on the following rules: (i) Identical CDR3 sequences with different V/J genes will be merged and only keep the V/J gene with the highest frequency; (ii) Different alleles of the same V/J family will be merged and only keep the family information; (iii) Only sequences including the V gene, J gene, and entire in-frame CDR3 sequences will be kept; (iv) The whole CDR3 sequence should begin with the cysteine (C), finish with the phenylalanine (F) or tryptophan (W), and contain no stop codon according to the IMGT (ImMunoGeneTics) guidelines [18]; (v) Then the CDR3 sequences with lengths fewer than 8 amino acids (AA) and larger than 24 AA will be eliminated.

### Measure TCR repertoire diversity by Renyi entropy

The repertoire diversity quantifies the number of different T-cell clones and their frequencies. We employ the widely-used Renyi entropy to estimate the TCR repertoire diversity [19]. Renyi entropy can graphically reflect the distribution of abundant clones and rare clones within a specific repertoire and quantify repertoire diversity. Here *p*_i_ is the frequency of the sequence in the TCR repertoire; *N* is the number of unique sequences in the TCR repertoire; and *b* is the base of the logarithm, which dictates the choice of entropy measurement units. The value of *α* determines the sensitivity of the diversity index to the sequence abundance in the TCR repertoire, as illustrated in formula (1). When *α* − > 0, all sequences have equal weight. The measure of entropy becomes a function of the number of unique sequences, but not dependent on their abundance. When *α* − > 1, the Renyi entropy equates to the Shannon entropy.

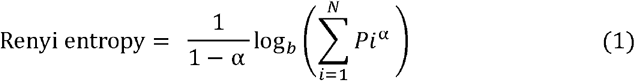

### Calculate TCR repertoire clonality by 1-Pielou’s index

The description of species abundance equivalence may also be used to quantify the dominance of clones in a TCR repertoire; this is known as clonal evenness [20]. The clonality score is the complement of clonal evenness score, which can be calculated using 1-Pielou’s index. As shown in formula (2), a clonality score of 0 indicates a maximally diverse population with even frequencies, whereas a value near to 1 indicates a repertoire driven by clonal dominance. Here, *pi* is the frequency of the sequence; *N* is the number of unique sequences, and *b* is the base of the logarithm.

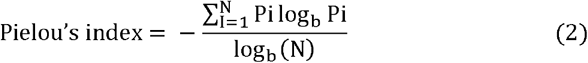

### Cluster CDR3 sequence and construct network

The similarity in CDR3 sequences implies structural resemblance for antigen recognition, which may result in shared antigen specificity [21]. Clustering similar CDR3 sequences is therefore essential for identifying cancer-associated receptors. GIANA is the best tool for clustering large-scale TCR repertoire (> 10^6^ sequences) with higher clustering accuracy, specificity and efficiency [15]. GIANA converts the sequence alignment and clustering problem into a classic nearest neighbor search in high-dimensional Euclidean space. We integrate the GIANA into network analysis in TCRosetta to cluster CDR3 sequences and then construct a network. The network analysis workflow contains the following steps: (i) Use GIANA to cluster similar CDR3 sequences; (ii) Use the Muscle [22] to align sequences obtained in the previous step, and then convert it to a distance matrix by computing Hamming distance between these sequences; (iii) Convert the distance matrix into an adjacency matrix in which two sequences are linked if their Hamming distance is less than 3; (iv) Construct a network using igraph (https://igraph.org/) based on the distance matrix from the previous step; (v) Calculate the betweenness of each network node using betweenness centrality, which is a measure of the centrality in a graph based on the shortest paths and represents the control of the node in graph theory; (vi) Combine the betweenness and degree as the weight of a node in the network; (vii) Use the random walk algorithm (*step* = 2) [23] to discover communities in the network; (viii) Make the sequence logo for each network communities using Logomaker [24]

### Calculate the public score of CDR3 sequences

V(D)J recombination creates diverse TCR sequence by assembling different germline V, D an J combinations and by diversifying junctions between these segments through nucleotide deletions and insertions. [25]. Several methods for calculating the probability of generation by V(D)J recombination have been developed. Here, the public score is defined as the generation probability of a CDR3 sequence in healthy individuals. OLGA [17] is the best approach for computing the generation probability of TRB CDR3 sequence, balancing the accuracy and efficiency through dynamic programming methods. We calculate the public score in TCRosetta using OLGA. The V(D)J recombination model for healthy individuals is obtained from the default human T cell beta chain model of OLGA using *–humanTRB* option. In addition, we constructed a background by computing the public score distribution for 5,000,000 CDR3 sequences randomly selected from collected healthy samples [26]. The Mann–Whitney U test is then used to examine the difference between the public score distributions of input data and the background in order to determine if the input data are healthy.

### Embedding analysis based on 3-mer CDR3 motif

Previous research indicated that only partial CDR3 sequences referred to as “motifs” contact specific peptides, which is a part of the TCR specificity [27]. We employ the k-mer (*k* = 3) abundance distribution to represent the feature of the TCR repertoire based on this discovery. Each TCR repertoire is represented as a 3-mer abundance distribution over 20^3^ = 8000 dimensions. The high dimension provides more particular information on TCR repertoire, but it also contributes lots of noise into the data. We applied PHATE [28] to embed TCR repertoire by producing a low-dimensional embedding feature vectors, which gives an accurate, denoised representation of a TCR repertoire explicit for visualization and is highly scalable in terms of memory and runtime. We trained a PHATE model to learn the embedding representation of k-mer abundance distribution of TCR repertoire using TCR data from eight diseases (Breast cancer, COVID-19, Crohn’s disease, Melanoma, Yellow fever vaccine, classical Hodgkin lymphoma, Non-small cell lung cancer and Cytomegalovirus). The following steps are performed to train the model. (i) For each sample, split all sequences into 3-mer using a sliding window with a step length of 1; (ii) Count all 3-mer motifs to form a TCR motif count matrix and exclude samples with either a large (top 20%) or a small number (bottom 20%) of motifs; (iii) Filter out motifs that do not appear in more than 50 percent of the disease samples, as a disease-specific motif should be present in majority of the samples of this disease; (iv) To exclude non-disease-specific motifs, we use the Mann–Whitney U test with P-value > 0.05 to calculate the difference between the distribution of motifs in healthy samples and disease samples; (v) Remove the batch effect by scprep (https://scprep.readthedocs.io/en/stable/index.html) and normalize the motif count matrix from eight diseases using Z-score; (vi) Use PHATE to train a distribution model.

### Annotation-based disease state statistics

Available TCR sequences with clinical information inspires us to investigate the possibility of annotating clinical information for unknown samples, a function that is absent from all existing web server. The batch search and annotation function of TCRosetta enables users to search multiple CDR3 sequences in our reference consisting of 244 million high-quality and annotated CDR3 sequences spanning 55 clinical diseases. We use Elasticsearch (https://www.elastic.co/), a fast and distributed search engine, to fuzzy search similar CDR3 sequences with a maximum of 1 mismatch in reference. As just a few CDR3 sequences are potential cancer-associated sequences and these specifical sequences are often cloned at high frequency in a TCR repertoire [29]. TCRosetta only keeps the top 3000 CDR3 sequences (ordered by frequency) for annotation. On the basis of the annotation results, we then access potential disease states using statistics methods. Sequences annotated for different diseases are more likely non-specific sequences, hence we exclude sequences annotated with more than five various diseases in the annotation results. To further investigate the possible disease state for the upload data, we use the Fisher Exact test to calculate *pi* for disease *i*, which measures the significance of the difference between the input data and the reference, as shown in formula (3). Only diseases with *pi* < 0.05 will be left. We calculate the *Mi* for each disease *i*, which represents the number of sequences annotated with this disease. Here, *N* is the total number of sequences in annotation results, *Z* is total number of sequences in reference. *Si* is the number of sequences in reference for disease *i*.

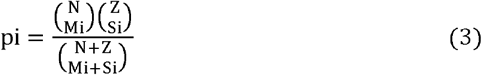

### Predict TCR-peptide binding

TCR recognition of antigen-specific peptides bound to MHC molecules. Predicting the binding scores between TCR and peptides is pivotal for development of immunotherapy and virus vaccines. We predict the binding scores between TRB sequences and peptides using ERGO-II. ERGO-II employs natural language processing (NPL) in the prediction model and uses the CDR3 alpha segment, the CDR3 beta segment, the MHC typing, and the V and J genes as the feature of predictor. It provides high accuracy prediction of TCR-peptide for previously unseen peptides [30]. We additionally assemble a peptide pool of 73 peptides by searching high-quality (confidence score = 3, TRB, HomoSapiens) peptides from VDJdb [31].

### Data availability

TCR repertoire sequencing data of both lung cancer and anti-PD-1 immunotherapy for oral carcinoma were downloaded from Adaptive Biotechnologies immuneACCESS online database (https://clients.adaptivebiotech.com/immuneaccess). All TCR-Seq datasets used in annotation analysis were obtained from public data repositories, with detailed information at http://bioinfo.life.hust.edu.cn/TCRosetta/#/document. The 73 high-quality peptides obtained from the VDJdb (https://vdjdb.cdr3.net/). The source code of TCRosetta is available at (https://github.com/yttyhhh/TCRosetta_code).

## Results

### TCRosetta overview

TCRosetta is a robust server for analyzing and visualizing the TCR repertoire. Because the TRB is more diverse than the TRA, the majority of TCR-Seq data and research focuses solely on the TRB [27]. Thus, all TCR repertoire analyses in TCRosetta pertain to the TRB. TCRosetta offers a user-friendly file manager for users to upload and manage data (upload, remove, and clear). It accepts four different types of input data: (i) Output file of V(D)J junction mapping software such as MiXCR [32], IMSEQ [33], CATT [34] or RTCR [35]; (ii) CDR3 sequences; (iii) AIRR-compatible formatted file and (iv) CSV formatted file. TCRosetta performs analyses depending on users’ selection in order to accommodate a variety of user requirements and decrease time expenditure. The input data will undergo pre-processing to guarantee data quality. TCRosetta provides general and advanced analyses of TCR repertoire. The general analyses include repertoire diversity, CDR3 length distribution, V/J gene usage, V-J gene utilization, motif, clonal composition and clonality. The advanced analyses contain network analysis, public analysis, embedding analysis, TCR-peptide binding prediction and enrichment analysis (Figure 1).

**Figure 1.**
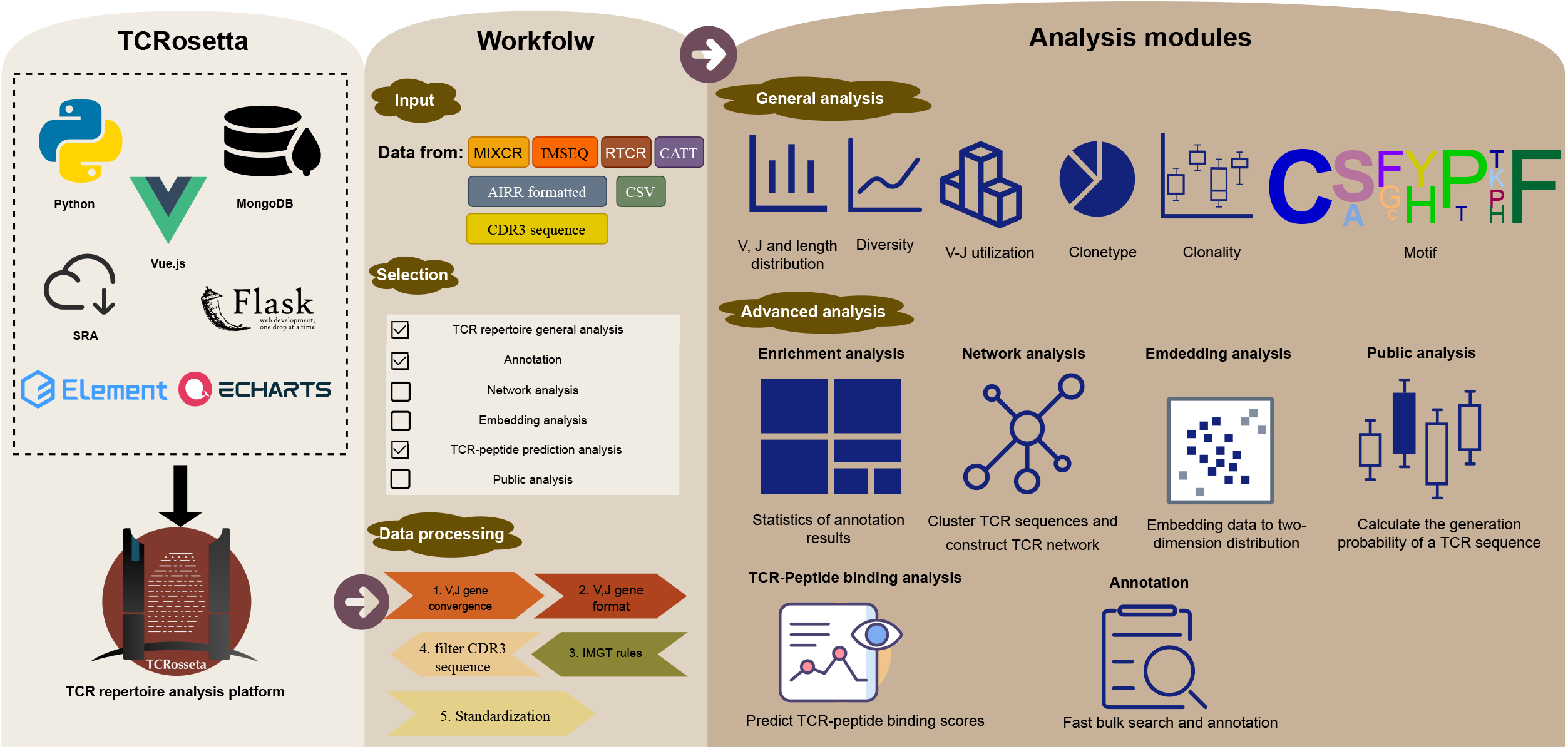
Overview of TCRosetta. For the website front end, we use Vue.js and echarts to build the web server and communicate with user clients. For the back end, we build the web server using Flask and Python 3.7. TCRosetta supports several input formats. Users can select different analysis modules. Data pre-processing is used to obtain reliable and high-quality TCR sequences. TCRosetta contains general analysis and advanced analysis.

### Comparison of TCRosetta with other existing TCR data analysis tools

Rapid expansion of high-throughput sequencing studies on T cells has led to the development of specific tools for TCR data analyzing. The downstream analysis of TCR data is often performed by different tools requiring diverse operating environments and expertise. For example, a suite of tools is often used to do TCR analysis. TCR general analysis may be performed by the tcR [10] package, generation probability is calculated by OLGA [17], similar TCR is clustered by GIANA [15], and TCR-peptide binding may be predicted by ERGO-II [30]. Although some integrated web servers and stand-alone tools have been developed to solve this problem, the majority of these tools are only designed to analyze the general features of TCR data. The restrictive format of their built-in TCR data makes it difficult for users to analyze according to their own data format. Importantly, all of these web servers require registration and login (details in Table 1).

**Table 1.** Comparison of TCRosetta with other existing TCR data analysis tools

| Server\Software\Package | TCRosetta | VDJserver | Vidjil | VDJviz | ImmunoSEQ analyzer | VDJtools | ImmunExplorer | immunarch | tcR |
| --- | --- | --- | --- | --- | --- | --- | --- | --- | --- |
| Type | server | server | server | server | software | software | software | R package | R package |
| Available | free | register | register | register | commercial | free | free | free | no longer supported |
| **CDR3 length distribution** | √ | √ | - | √ | √ | √ | - | √ | √ |
| **Diversity** | √ | √ | √ | √ | - | √ | √ | √ | √ |
| **Clonality** | √ | √ | - | √ | - | √ | √ | √ | √ |
| **V/J gene usage** | √ | √ | √ | √ | √ | √ | √ | √ | √ |
| **V-J gene utilization** | √ | - | - | √ | √ | √ | - | √ | √ |
| **Clonal composition** | √ | - | - | √ | √ | √ | √ | √ | √ |
| **Sequence logo** | √ | - | - | - | - | - | - | - | √ |
| **Annotation** | √ | - | - | - | - | √ | - | √ | - |
| **Enrichment analysis** | √ | - | - | - | - | - | - | - | - |
| **Cluster similar sequences** | √ | - | - | - | - | √ | - | - | √ |
| **Calculate generation probability** | √ | - | - | - | - | - | - | - | √ |
| **Network analysis** | √ | - | - | - | - | - | - | - | - |
| **TCR-peptide binding prediction** | √ | - | - | - | - | - | - | - | - |
| **Embedding analysis** | √ | - | - | - | - | - | - | √ | √ |

The only three web servers for TCR analysis, VDJserver [36], Vidjil [37] and VDJviz [38], all require registration and can only do general features analysis of repertoire (diversity, V/J gene usage, CDR3 length distribution) with no additional advanced analysis. The workflow of VDJserver is complex including account creation, project creation, file uploading, file attributes setting and analysis. The R package, tcR [10], can do some general and advanced analysis, however it is no longer supported. ImmunoSEQ analyzer (https://www.immunoseq.com/analyzer/) is a non-free commercial analytics platform included in the immunoSEQ Service. ImmunExplorer [39] is a software for only analyzing diversity and clonality of TCRs. VDJtools [11] and Immunarch [40] are immune repertoire exploration tools. These two tools do not provide the latest TCR analysis methods like network analysis, TCR-peptide binding prediction and TCR sequence generation probability.

TCRosetta is a one-stop solution for TCR data analysis that interacts with users via a web-based GUI and does not require coding knowledge or command line experience. TCRosetta is superior to existing web servers, software and packages for TCR data analysis in following ways (Table 1): (i) It is the first web server to integrate the majority of known TCR data analysis methods. (ii) It enables the quick batch searches for similar TCR sequences and annotates disease information in large-scale reference data. (iii) It offers a user-friendly interface, interactive and downloadable graphs, and personalized analysis options.

### Analysis modules in TCRosetta

TCRosetta consists of six analysis modules: General, Annotation, Enrichment, Public, Embedding, Network, and TCR-peptide binding prediction (Figure 2). It is a web server for TCR analysis from many aspects.

**Figure 2.**
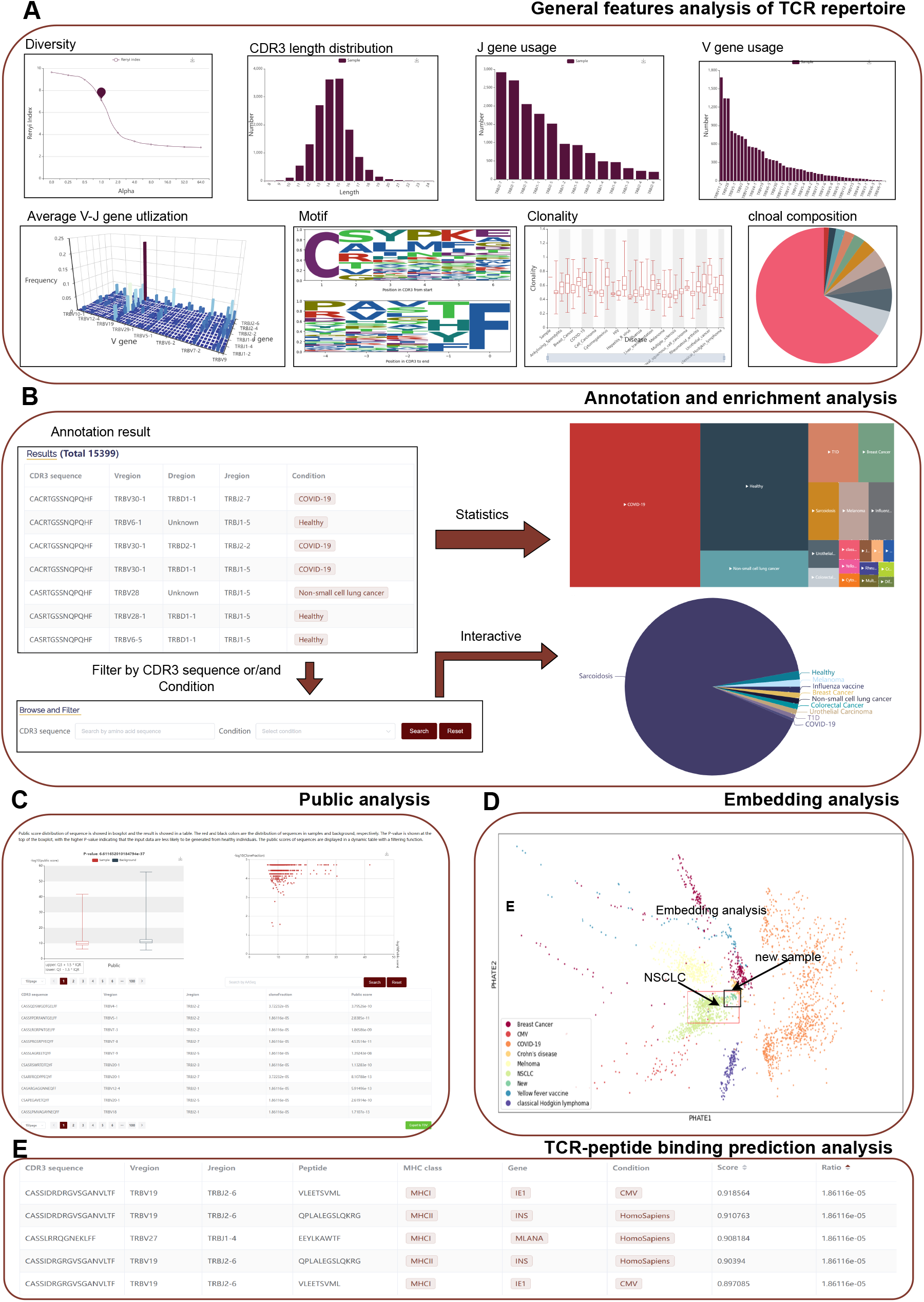
Analysis modules of TCRosetta. (A) General analysis of TCR repertoire. The TCR repertoire features of the upload file include V/J gene usage, diversity, clonality, CDR3 length distribution and clonal composition. (B) Annotation and enrichment analysis. The enriched disease distribution of upload data is displayed in a treemap. Different square colors represent different diseases, and the size of the square is the number of unique sequences annotated with this disease. Users can further browse the CDR3 sequence, V and J usage by clicking the square. The corrected enriched disease distribution is shown in a pie chart. The size of the sector represents the ratio of the disease. (C) Public analysis. The public score distribution of the sample is displayed in an interactive box chart. The red and black colors are the distribution of sequences in the ample and reference, respectively. The P-value is shown at the top of the boxplot. The relationship between public score and frequency of CDR3s in TCR repertoire is displayed in a scatter chart. The public scores of sequences are displayed in a dynamic table with a filtering function. (D) Embedding analysis. The result of embedding analysis on an independent test set. The legend is at the bottom left of the picture, and “New” in the legend represents the samples in the test set. The green points in the red box result from embedding analysis for samples in the test set. (E) TCR-peptide binding prediction analysis. The prediction results are displayed in a dynamic table.

### The General module

Some changes of TCR repertoire, such as the diversity and clonality, enable sensitive tracking of dynamic changes in antigen-specific T cells [41]. The General module performs general TCR repertoire feature analysis (Figure 2A) including: (i) the CDR3 length distribution; (ii) the diversity; (iii) the V-J gene utilization; (iv) the sequence logo of the first and last five amino acids of CDR3 sequences; (v) the V/J gene usage; (vi) the clonality and its’ distribution; (vii) the clonal composition. The general analysis results are displayed on the “General analysis of TCR repertoire” page.

### The Annotation and Enrichment modules

TCRosetta offers Annotation and Enrichment modules to annotate potential disease states for unknown TCR sample (Figure 2B). These modules annotate sample by batch searching similar sequences in manually curated disease-related TCR database and then using the search result to calculate the disease enrichment score in sample. Results of annotation are displayed in a dynamic table with filter and export function on the “Annotation by batch search” page. The enrichment results are displayed in the “Enrichment analysis” page. The enriched disease distribution of upload data is displayed in a treemap. Different square colors represent different diseases, and the size of the square is the number of unique sequences annotated with this disease. The corrected enriched disease distribution is shown in a pie chart.

### The Public module

Due to the high diversity of the TCR, the majority of TCRs are private to each individual and do not exist in other people. However, some TCRs are public and shared by multiple individuals. TCRosetta calculates the public score of CDR3s and obtains the TCR repertoire public score distribution using OLGA [18] in the public module. To detect if the public score distribution of input samples is skewed, TCRosetta additionally computes the distributional difference between sample and reference. The public analysis results are displayed on the “Public analysis” page (Figure 2C). The public score distribution of the sample is displayed in an interactive box chart. The P-value is shown at the top of the boxplot, with the higher P-value indicating that the input data are less likely to be generated from healthy individuals. The relationship between public score and frequency of sequences in TCR repertoire is displayed in a scatter chart. The public scores of sequences are displayed in a dynamic table with a filtering function.

### The Embedding module

CDR3 is the primary region of TCR that contacts peptides, and the distribution of CDR3 sequence motifs may reflect the disease-specific information. The embedding module splits all CDR3 amino acid sequences in uploaded data into k-mer to uncover possible disease specificity from CDR3 motif distribution and obtain a high-dimensional distribution of motifs. Then, TCRosetta embeds it into a model pre-trained by PHATE [28]. To evaluate the performance of the embedding analysis, we conducted an independent validation with 38 TCR-seq samples from 16 lung cancer patients. We embedded the data set into two-dimensional space by our pre-trained model and found that most points of the data set fell in the region of Non-small Cell Lung Cancer (NSCLC) samples (Figure 2D). The embedding analysis results are displayed on the “Embedding analysis” page

### The TCR-peptide binding prediction module

Many TCR responses have high level of cross-reactivity, with a single TCR binding a large number of peptides and a single MHC-peptide binding a large number of TCRs. TCRs that bind the same MHC-peptide may share similarities. The prediction module predicts binding scores between TCRs and peptides (Figure 2E). The prediction results are displayed in a dynamic table with filtering and exporting functions on the “TCR-peptide prediction” page.

### The Network module

Networks have been used to show immune responses defined by the similarity between sequences [42]. Such as, network connection was also used to distinguish clonally expanded repertoires from individuals with HIV-1 infection [43] and chronic lymphocytic leukemia [44]. The Network module transforms a TCR repertoire into a TCR network by GIANA and iGraph, and then divides the network into different communities (Figure 3A). Each community consists of similar CDR3 sequences and the sequence similarity within a community is greater than that of outside the community. The clustering results are displayed in a dynamic table with filtering and exporting functions (Figure 3B). To better observe the usage of amino acids in each position of sequences in each community, we make the sequence logo to show the position weight matrix of the complete CDR3 sequence, reflecting community conservation (Figure 3C). The network, sequence logo and cluster result are shown on the “Network analysis” page.

**Figure 3.**
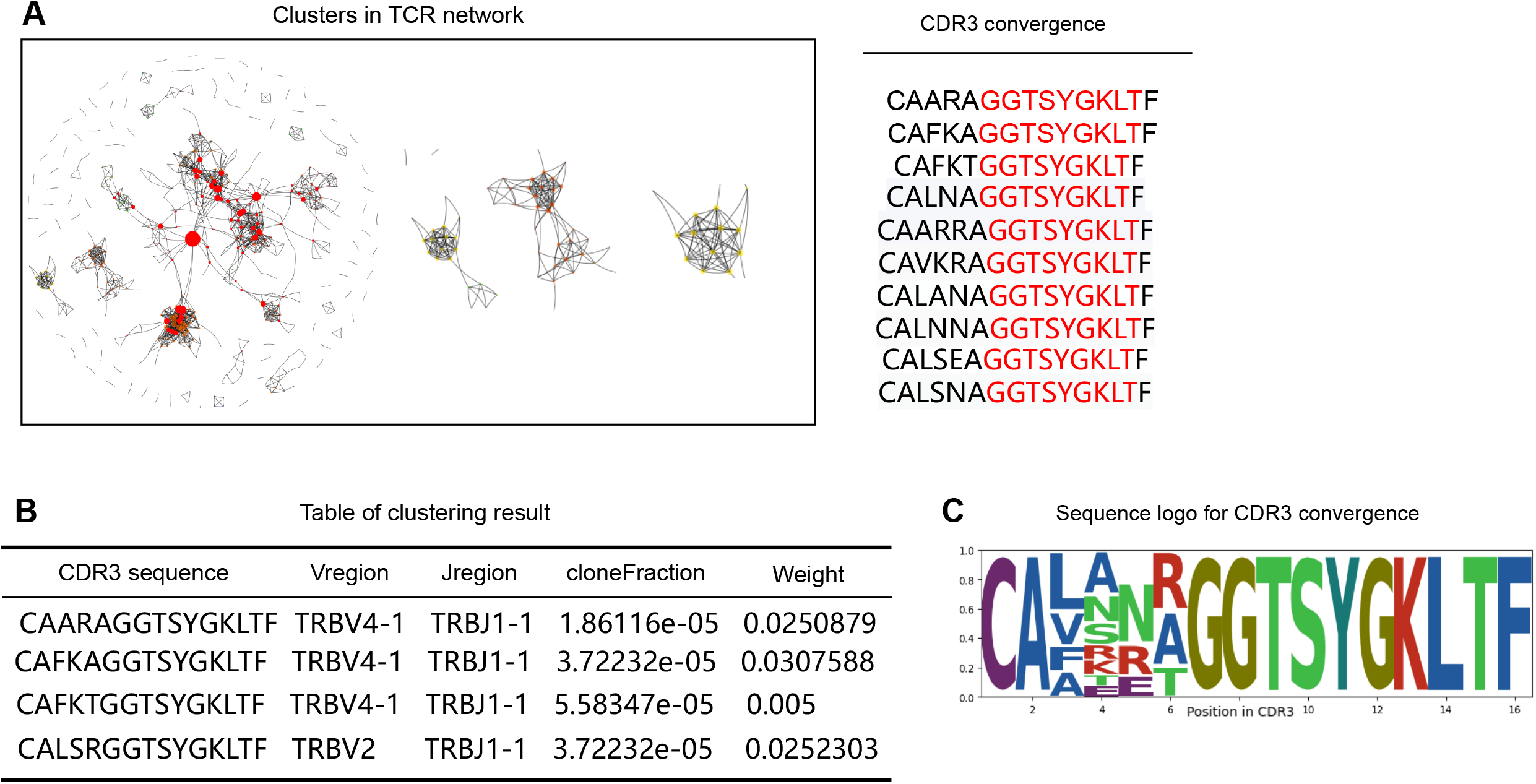
Results of network analysis in TCRosetta. (A) Network of TCR sequences. The size of nodes represents the weight of this sequence. Different color represents different communities. The red part of CDR3 sequences is the conserved residue in this community. (B) Table of the clustering result. Each column contains CDR3 sequence, Vregion, Jregion, cloneFraction, and Weight. (C) Sequence logo of CDR3 convergence in the network.

### Case Study of TCRosetta Application to Identify Different Oral Cancer Characteristics Before/After Anti-PD-1 Immunotherapy

Ten pre-treatment and ten post-treatment samples from an independent dataset of anti-PD-1 immunotherapy for oral carcinoma [29] were used to illustrate the powerful function of TCRosetta in TCR repertoire analysis. Pre-treatment and post-treatment samples yielded 136,992 and 174,885 unique TRB CDR3 sequences containing the V/J gene information, respectively. Two groups were separately uploaded into TCRosetta with all analyses selected. The results are summarized in Figure 4. By comparing the diversity curves of two groups, we found the curve dropped more quickly after treatment, suggesting that there were more abundant clones in treated group (Figure 4A). It may demonstrate that anti-PD1 therapy can result in the expansion of some T cells [45]. We also found the clonality increased (pre-treatment:0.501, post-treatment:0.694) during the treatment, which is consistent with the original study’s finding. In the post-treatment, the CDR3 sequence with the highest clonal frequency (CATSRESPGQGIDEQ) differs from the pre-treatment group (CASSEEAGTIYEQYF) (Figure 4B). TRBV5-1/TRBJ1-2 is the most frequently used V-J gene pairing segment in the pre-treatment group, while the TRBV15-1/TRBJ2-1is the most frequently used segment in the post-treatment group (Figure 4C). Some V and J gene segments, such as TRBV7-9 and TRBV2-1, grow during treatment (Figure 4D). It is likely that the treatment enhanced the activation and expansion of T cells, resulting in an increased abundance of these V/J genes or CDR3s in the tumor. By clustering similar CDR3 sequences in network analysis module for the pre-treatment and post-treatment groups, we studied the potential cancer-associated CDR3 sequences. We found that the distributions of network nodes in the two groups are different (Figure 4E). By analyzing each cluster in network, we discovered that “GTG” are the conserved residues based on the amino acid usage frequency in the biggest cluster of the pre-treatment group, which differs from post-treatment group and may be a favorable residue for antigen binding (Figure 4E). To investigate further if there is a difference in binding peptides of these CDR3s in network, we predicted TCR-peptide binding for pre-treatment and post-treatment groups. We found that the TCR repertoire could bind more peptides after treatment, with 9 peptides with binding scores > 0.7 in the pre-treatment group and 33 in the post-treatment group (Figure 4F).

**Figure 4.**
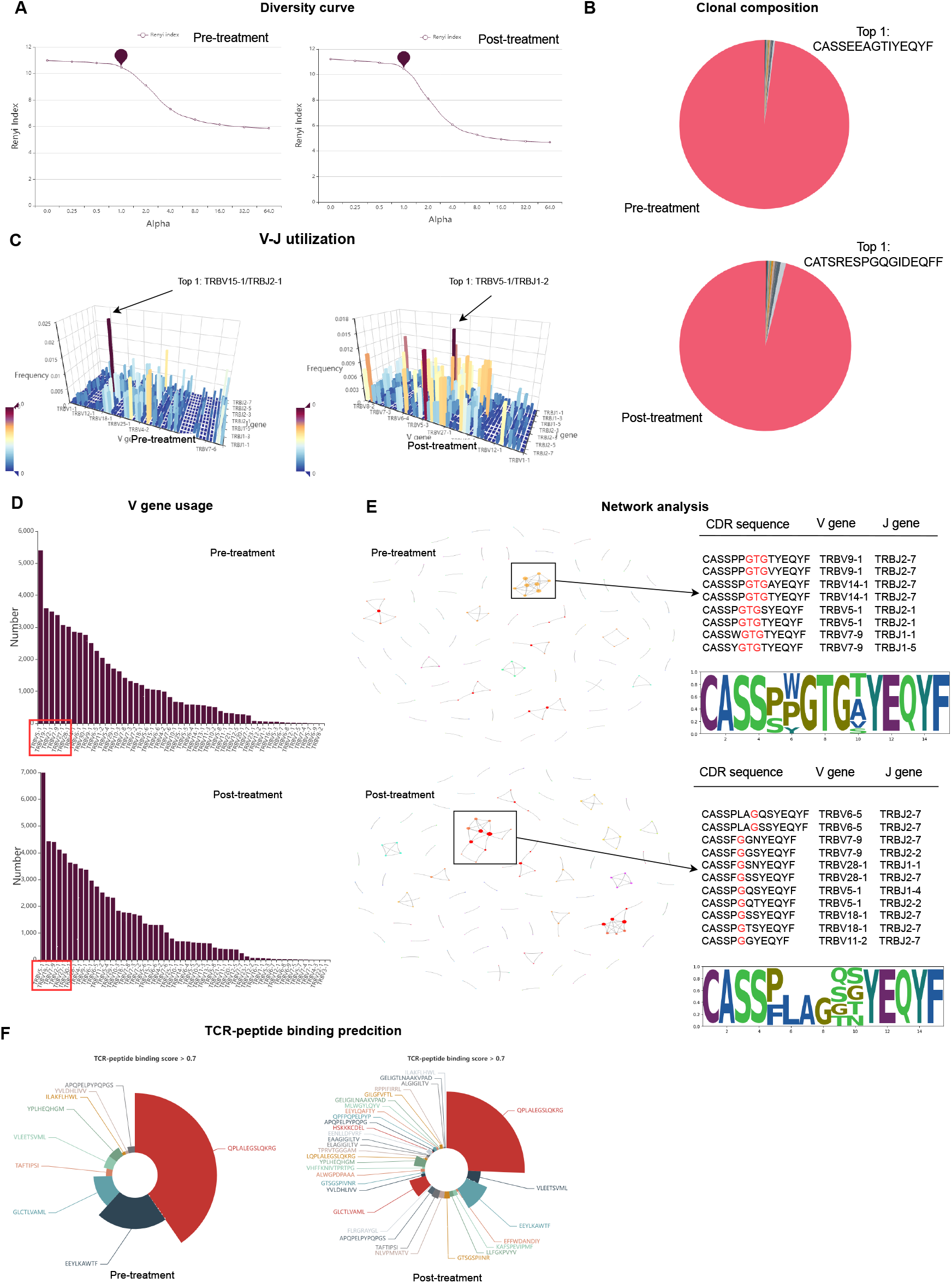
A case study is analyzing T-cell receptor repertoire by TCRosetta. (A) The Renyi diversity curve of the pre-treatment group and the post-treatment group. The slope gradient increases as the repertoire’s distribution become more monoclonal. (B) The TRB CDR3 clone type distribution in the pre-treatment and post-treatment groups. The top1 clone is shown in the figure. (C) The TRB V-J gene pairing usage of two groups. The color represents the frequency, whereas the darker color represents higher V-J gene pairing usage. The top1 V-J gene pairing usage is shown in the figure. (D) The TRB V gene usage in two groups. (E) The network and sequence logo of the pre-treatment group and the post-treatment group. TRB CDR3 sequences and the sequence logo of the cluster in the network (black box) are shown on network’s right. Conserved CDR3 motif residues in the middle are highlighted in red in CDR3 alignments. (F) TCR-peptide binding prediction analysis results. Different color in chart represents the different peptide. The size of the sector represents the ratio of the peptide.

## Discussion

The analysis of the TCR repertoire may help to gain a better understanding of the immune system. However, these analysis tools require extensive skills in bioinformatics and have different limitations. Thus, we developed TCRosetta, a comprehensive platform for analyzing T-cell repertoire. Not only could it analyze the features of TCR repertoire and display them in interactive graphs but it also the first platform with a batch search and TCR annotation function.

TCRosetta can be applied to many situations. For example, calculating the features (CDR3 sequence length distribution, diversity, V-J utilization, and clonality) of the TCR repertoire can be used as biomarkers for predicting response to immunotherapy. Besides, Network analysis could help to identify T-cell clones specific to the tumor by clustering similar TCR sequences, which could play an important role in some tumor immunotherapy and TCR-T therapy. Moreover, Public analysis of TCRosetta may assist in predicting the autoimmune disease prognosis by discovering TCR CDR3 sequences shared between individuals associated with self-related immunity.

As an attempt at a comprehensive interpretation of the TCR repertoire, our study has several limitations. First, TCRosetta mainly studies the features of the TCR beta chain and ignores the alpha chain. Although the TCR beta chain has the highest diversity and plays an important role in tumor antigen recognition, some studies have shown that the TCR alpha chain also has functions in tumor antigen recognition. Second, TCRosetta integrated more than 244 million reliable TRB CDR3 sequences with the clinical condition in annotation reference, which makes it possible to annotate TCR repertoire with big data. Due to the high diversity of TCRs in the human body (range from 10^12^ to 10^18^), these data may not be sufficient to annotate all TCRs. Third, users can not upload multiple samples at the same time for comparative analysis.

With the development of TCR studies, we will continue to add more analysis functions for the TCR repertoire analysis and support more TCR profiling tools. We will further improve the data volume of the reference to ensure that more TCR sequences can be annotated and allow users to upload multiple samples for analysis. We believe that TCRosetta would facilitate TCR-related research and clinical applications.

## Acknowledgements

Not applicable

## List of Abbreviations

TCR: T-cell receptor
CDR3: Complementary determining region 3
MHC: Major histocompatibility complex
TRB: TCR beta chain
AA: Amino acid
V: Variable genes
J: Joining genes
D: Diversity genes
C: Cysteine
F: Phenylalanine
W: Tryptophan

